# Discrimination of inherent characteristics of susceptible and resistant strains of *Anopheles gambiae* by explainable Artificial Intelligence Analysis of Flight Trajectories

**DOI:** 10.1101/2024.12.10.627548

**Authors:** Yasser M. Qureshi, Vitaly Voloshin, Katherine Gleave, Hilary Ranson, Philip J. McCall, Cathy E. Towers, James A. Covington, David P. Towers

**Affiliations:** School of Engineering, University of Warwick, Coventry, CV4 7AL, UK; School of Biological and Behavioural Sciences, Queen Mary University of London, London, E1 4NS, UK; Vector Biology Department, Liverpool School of Tropical Medicine, Pembroke Place, Liverpool, L3 5QA, UK

## Abstract

Understanding mosquito behaviours is vital for development of insecticide-treated bednets (ITNs), which have been successfully deployed in sub-Saharan Africa to reduce disease transmission, particularly malaria. However, rising insecticide resistance (IR) among mosquito populations, owing to genetic and behavioural changes, poses a significant challenge. We present a machine learning pipeline that successfully distinguishes between IR and insecticide-susceptible (IS) mosquito behaviours by analysing trajectory data. Data driven methods are introduced to accommodate common tracking system shortcomings that occur due to mosquito positions being occluded by the bednet or other objects. Trajectories, obtained from room-scale tracking of two IR and two IS strains around a human-baited, untreated bednet, were analysed using features such as velocity, acceleration, and geometric descriptors. Using these features, an XGBoost model achieved a balanced accuracy of 0.743 and a ROC AUC of 0.813 in classifying IR from IS mosquitoes. SHAP analysis helped decipher that IR mosquitoes tend to fly slower with more directed flight paths and lower variability than IS—traits that are likely a fitness advantage by enhancing their ability to respond more quickly to bloodmeal cues. This approach provides valuable insights based on flight behaviour that can reveal the action of interventions and insecticides on mosquito physiology.

## 1 Introduction

Mosquito-borne diseases are responsible for around 1 million deaths every year (1). These include some of the deadliest viral and parasitic infections affecting humans, such as malaria, dengue, yellow fever, Zika and filariasis (2). The threat from malaria remains the most significant, with the WHO Africa region accounting for > 90% of cases and deaths globally (3). In the African region, the main preventative measure is the insecticide treated net (ITN), the use of which has risen from < 5% of households in 2000 to over 50% by 2015 (3). Although malaria deaths fell by approximately 55% (normalised figures per 100,000 population) in the same period, malaria incidence and mortality rates have largely plateaued since then (3). The emergence of resistance to the pyrethroid insecticide used on bednets and its spread across sub-Saharan Africa is believed to be at least partially responsible for the lack of progress in malaria vector elimination targets.

All ITNs currently recommended for use by WHO contain pyrethroid insecticides and resistance to this insecticide class in malaria vectors has rapidly spread since the scale up of ITN use, nearly 25 years ago. Resistance can be caused by physiological changes in the mosquito that reduce the amount of insecticide reaching the target site or reduce the binding to the target, or behavioural mechanisms that reduce contact with the insecticide (4). In the major African malaria vector, Anopheles gambiae, multiple mechanisms causing either behavioural or physiological resistance have been reported. Behavioural resistance can arise if changes in the vector population’s preferred biting time (night v day) or location (indoor v outdoor) evolve to reduce the likelihood of contact with insecticide on bednets. There have been several reports of shifts in biting patterns (5–7) following ITN scale up; higher rates of outdoor transmission since the year 2000 are thought to have resulted in millions of additional malaria cases each year (8). Nonetheless the majority of mosquito bites can still be prevented by ITN use. Physiological mechanisms such as mutations in the sodium channel target, or increased rates of detoxification by cytochrome P450s can cause very high levels of pyrethroid resistance with other, less well-defined mechanisms, further contributing to the phenotype (9). The intensity of pyrethroid resistance, and the underlying mechanisms responsible differs between populations, sometimes even within a small geographical area (10).

The link between physiological resistance mechanisms and mosquito behaviour is poorly understood. Fitness effects of specific resistance mechanisms that impact their interaction with ITNs have been reported (11) but limitations in bioassay methods and the ability to measure mosquito behaviour have constrained understanding of the relationship between pyrethroid resistance and the mosquito’s interaction with ITNs.

More detailed analyses of vector behaviour has been achieved via video based optical tracking systems including behaviour of mosquitoes at human baited bednets in sub-Saharan Africa (12). That study determined that net contact of less than 1 minute per mosquito was sufficient to reduce activity of pyrethroid susceptible mosquitoes to a negligible level 30 minutes after the mosquitoes were introduced. Insights were gained regarding the preferred location of activity - above the bednet – and direction of arrival – descent from above onto the host from above.

Specific mosquito tracking analyses have used video tracking to compare flight activity of pyrethroid insecticide susceptible (IS; Ngousso and the highly susceptible Kisumu strains) and resistant (IR; VK7 and Banfora strains) strains of *Anopheles gambiae* as they respond to human hosts at different bednets: untreated (UT), Olyset (OL, single pyrethroid active ingredient), Permanet 3 (P3, pyrethroid on all net surfaces, roof with pyrethroid and piperonyl butoxide, PBO) and Interceptor G2 (IG2, pyrethroid and chlorfenapyr on all net surfaces) bednets (13). Significantly higher levels of activity were seen around an untreated net in comparison to the 3 ITNs, at which the total activity, the number and duration of net contacts were similar at all ITNs for both susceptible and resistant strains. There was a steep decay in mosquito activity after approximately 20 minutes, indicative of knockdown or mortality, for the IS strains at OL and P3 ITNs but only with the highly susceptible Kisumu at IG2. A slow decline in activity was measured with OL, P3 and IG2 nets and the IR strains indicating the slower impact of the second insecticide on the IR strains as they are resistant to the fast acting ‘knockdown’ effect of the pyrethroid.

In this study we investigated whether a more detailed examination of mosquito flight, using machine learning models, could identify basic or inherent behaviours that distinguish IS and IR populations, prior to insecticide contact. Explainable AI (XAI) is an emerging field that attempts to interpret machine learning models for better transparency and understanding. By uncovering the rationales behind decision-making, XAI can raise confidence in AI systems and enhance their real-world adoption. For example, Ryo et al successfully employed XAI to interpret ML species distribution models boosting the usability and interpretability of the ecological models for further research (13). That study demonstrated the potential for XAI to assist ecologists identify the underlying factors, connecting the key features that influence species distribution within an ML model.

Here, we applied XAI techniques to detect fundamental differences between insecticide-susceptible (IS) and insecticide-resistant (IR) mosquito strains using trajectory features. This study utilised trajectory data of malaria vectors orienting to a human host within an untreated bednet in order to examine behaviours without any effect of insecticides, i.e. the innate behaviours and inherent flight characteristics of pyrethroid resistant and susceptible strains of primary malaria vector *Anopheles gambiae s.l.*. This work has the potential to improve our understanding of how insecticide resistance impacts mosquito behaviour.

## 2 Results

### 2.1 Data Processing

Trajectories were split into segments to mitigate the impact of duration imbalance between tracks. Through hyperparameter tuning, various window sizes and overlap lengths for the windowing technique were tested. Each model performed best with different window parameters. Table 1 displays the window parameters selected after tuning and the effect they have on the number of tracks (tracks shorter than the segment length are discarded) and segments within the dataset partitioned for modelling. Note that the numbers of tracks and segments were after segment quality filtering.

**Table 1:** Windowing parameters for each model alongside the final number of tracks and segments.

| Machine Learning Model | Window Size (s) | Window Overlap (s) | Final Number of tracks | Number of segments |
| --- | --- | --- | --- | --- |
| IR vs IS, XGBoost | 8.0 | 7.5 | 10,514 | 695,509 |
| IR vs IS, Random Forests | 7.5 | 6.5 | 10,943 | 357,120 |
| IR vs IS, Logistic Regression | 9.5 | 9.0 | 9,589 | 664,369 |
| Kisumu vs N’goussu, XGBoost | 1.5 | 0.5 | 11,402 | 192,150 |
| Banfora vs VK7, XGBoost | 6.0 | 6.5 | 6,017 | 352,761 |
| Multiclass, XGBoost | 6.0 | 5.5 | 11,887 | 683,967 |

### 2.2 Model Evaluation - Classification of Insecticide Resistance Status

The performance of the classification of insecticide resistance status, IR or IS, is provided in Table 2. The average and the range (minimum and maximum) of scores across all folds are provided in brackets.

**Table 2:** Performance of each machine learning model applied to independent test data for IR (Banfora & VK7) vs IS (Kisumu & N’goussu), with the average and the range provided across all folds.

| Model | Balanced Accuracy | ROC AUC Score |
| --- | --- | --- |
| XGBoost | 0.743 (0.688 - 0.786) | 0.813 (0.757 - 0.866) |
| Random Forests | 0.721 (0.680 - 0.772) | 0.806 (0.760 - 0.859) |
| Logistic Regression | 0.723 (0.682 - 0.759) | 0.788 (0.734 - 0.831) |

The confusion matrix for each model is displayed in Figure 1 which shows the percentage of the predictions over all folds, and the ROC curves are depicted in Figure 2 featuring each folds’ curve. Additionally, a more detailed review of the performances of each tuned model is given in Table 3.

**Fig. 1:**
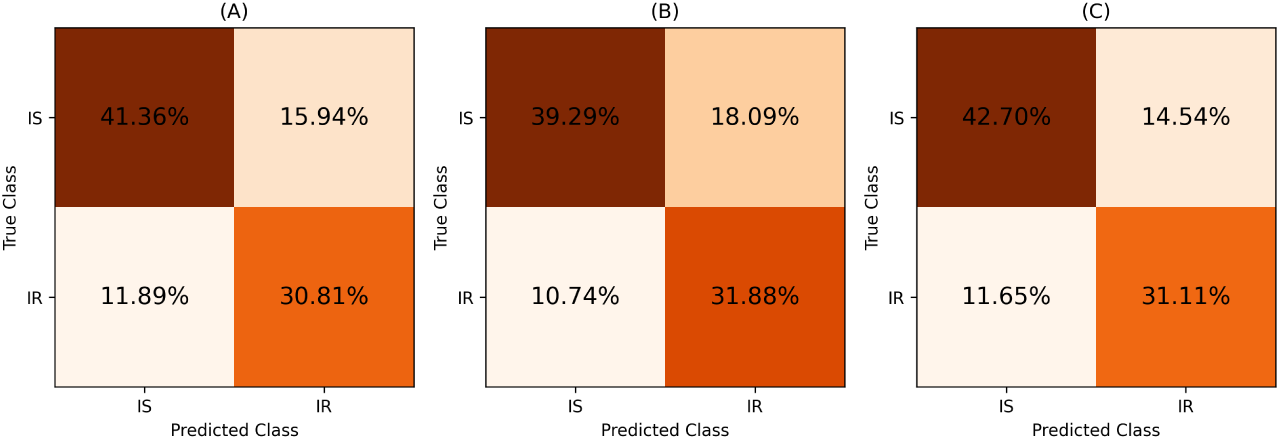
Confusion matrices from independent test data displaying the percentages of the predictions over all folds for IR vs IS. In the figure, (A) displays the logistic regression model, (B) is the random forests model and (C) is the XGBoost model.

**Fig. 2:**
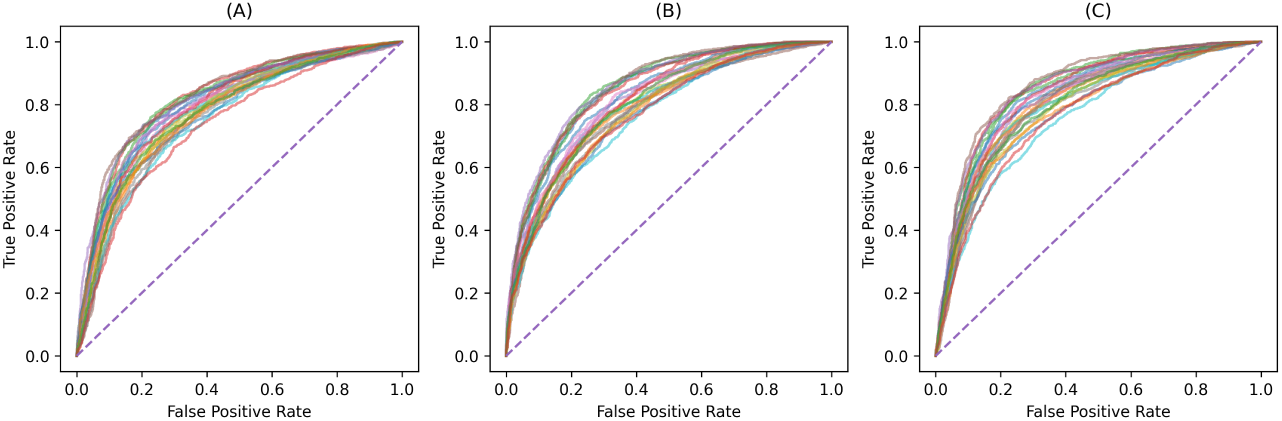
Confusion matrices from independent test data displaying the percentages of the predictions over all folds for IR vs IS. In the figure, (A) displays the logistic regression model, (B) is the random forests model and (C) is the XGBoost model.

**Table 3:** Performance metrics of each IR vs IS machine learning model applied to independent test data.

| Performance metric | Logistic Regression | Random Forests | XGBoost |
| --- | --- | --- | --- |
| F1 score (IR) | 0.686 (0.642 - 0.739) | 0.687 (0.638 - 0.747) | 0.702 (0.646 - 0.757) |
| F1 score (IS) | 0.746 (0.675 - 0.797) | 0.730 (0.658 - 0.776) | 0.765 (0.725 - 0.802) |
| Recall (IR) | 0.725 (0.631 - 0.798) | 0.758 (0.657 - 0.888) | 0.739 (0.628 - 0.883) |
| Recall (IS) | 0.720 (0.591 - 0.779) | 0.684 (0.543 - 0.780) | 0.746 (0.626 - 0.852) |
| Precision (IR) | 0.656 (0.540 - 0.750) | 0.637 (0.522 - 0.746) | 0.682 (0.574 - 0.801) |
| Precision (IS) | 0.780 (0.706 - 0.869) | 0.792 (0.698 - 0.914) | 0.794 (0.694 - 0.918) |
| PR AUC (IR) | 0.714 (0.621 - 0.788) | 0.750 (0.693 - 0.800) | 0.735 (0.681 - 0.793) |
| PR AUC (IS) | 0.812 (0.747 - 0.883) | 0.841 (0.779 - 0.919) | 0.840 (0.777 - 0.917) |
| Cohen Kappa Coefficient | 0.435 (0.351 - 0.508) | 0.425 (0.327 - 0.528) | 0.472 (0.373 - 0.556) |
| Matthew Correlation Coefficient | 0.440 (0.361 - 0.509) | 0.436 (0.356 - 0.538) | 0.480 (0.374 - 0.556) |
| Log loss | 0.567 (0.523 - 0.635) | 0.565 (0.539 - 0.605) | 0.552 (0.497 - 0.594) |

The SHAP plots for the best performing fold for the XGBoost model applied to independent test data are provided below. This includes SHAP summary plot, Figure 3, and SHAP bar plot, Figure 4.

**Fig. 3:**
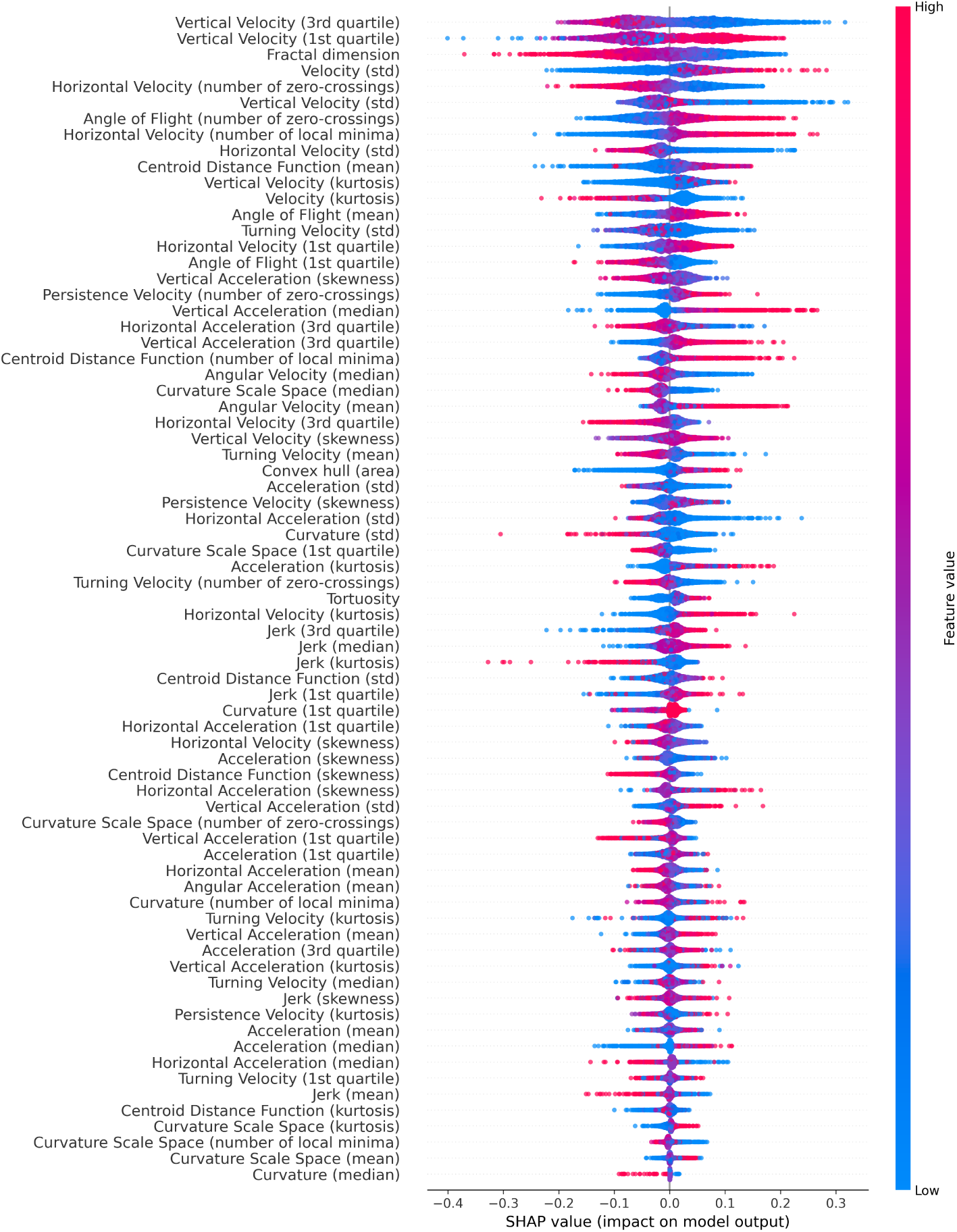
SHAP summary plot for the best IR vs IS XGBoost model fold applied to independent test data displaying the contributions of each model. Features are ordered by mean absolute SHAP value where each dot represents a segment, with its colour displaying its feature value. Positive SHAP values contribute towards the IR class, whilst negative SHAP values contribute towards the IS class.

**Fig. 4:**
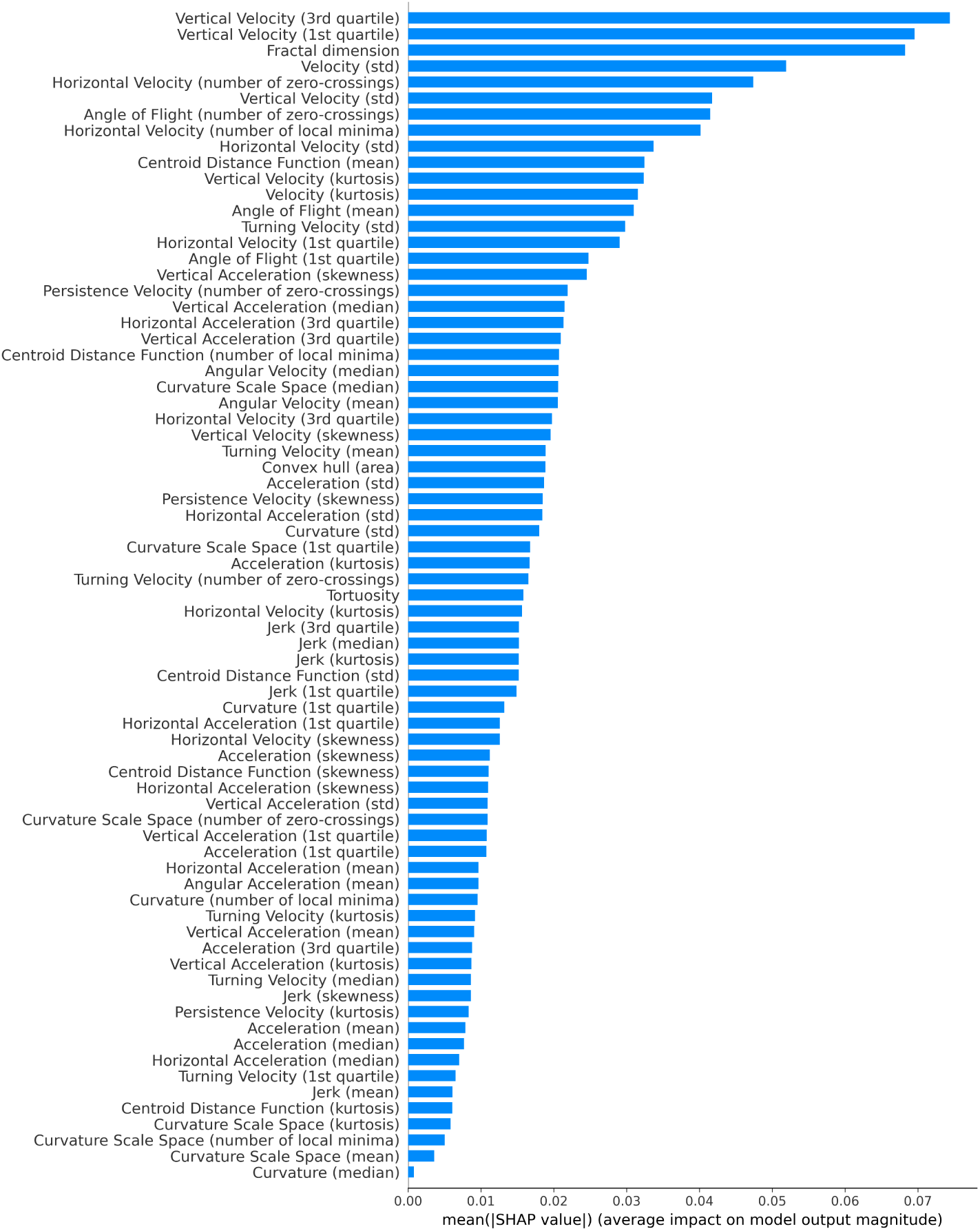
SHAP bar plot for the best IR vs IS XGBoost model fold applied to independent test data. The features are ordered by mean absolute SHAP value.

Velocity in the vertical direction is one of the strongest contributors to the model. There are 5 vertical-velocity features, ranked 1, 2, 6, 11 and 27 in the SHAPs bar plot. These are the 3rd quartile (identifying 75% of the population have lower values), 1st quartile (25%of the population), standard deviation, kurtosis and skewness respectively of the vertical-velocity distributions within a track segment. Figure 5 displays the SHAP scatter plots for these vertical veloc-ity features. A histogram comparison of vertical velocities for IR and IS, Figure 6, reveals distinct distributions that are consistent with the SHAP scatter plots.

**Fig. 5:**
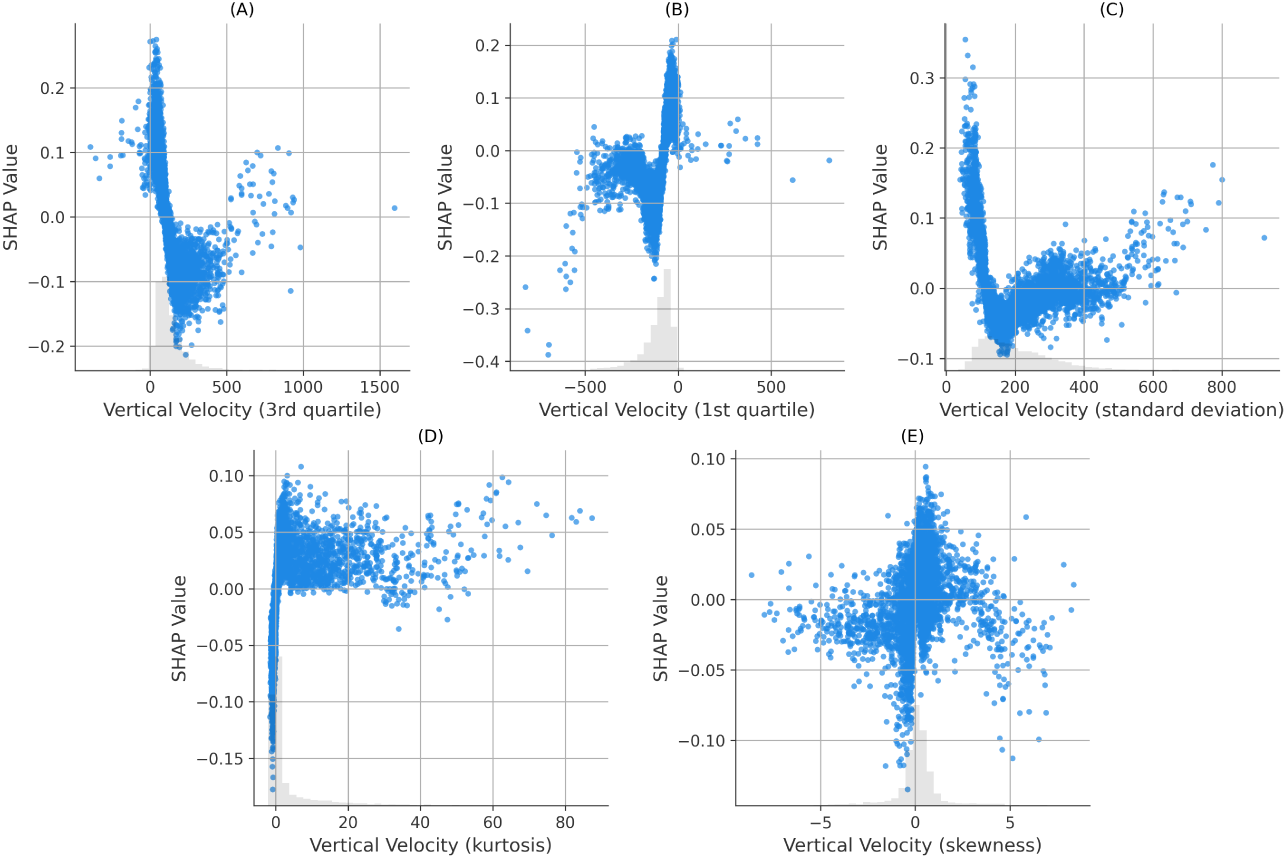
SHAP scatter plots for the vertical velocity features. (A) is the 3rd quartile of vertical velocity, (B) is the 1st quartile of vertical velocity, (C) is the standard deviation of vertical velocity, (D) is the kurtosis of vertical velocity, and (E) is the skewness of vertical velocity.

**Fig. 6:**
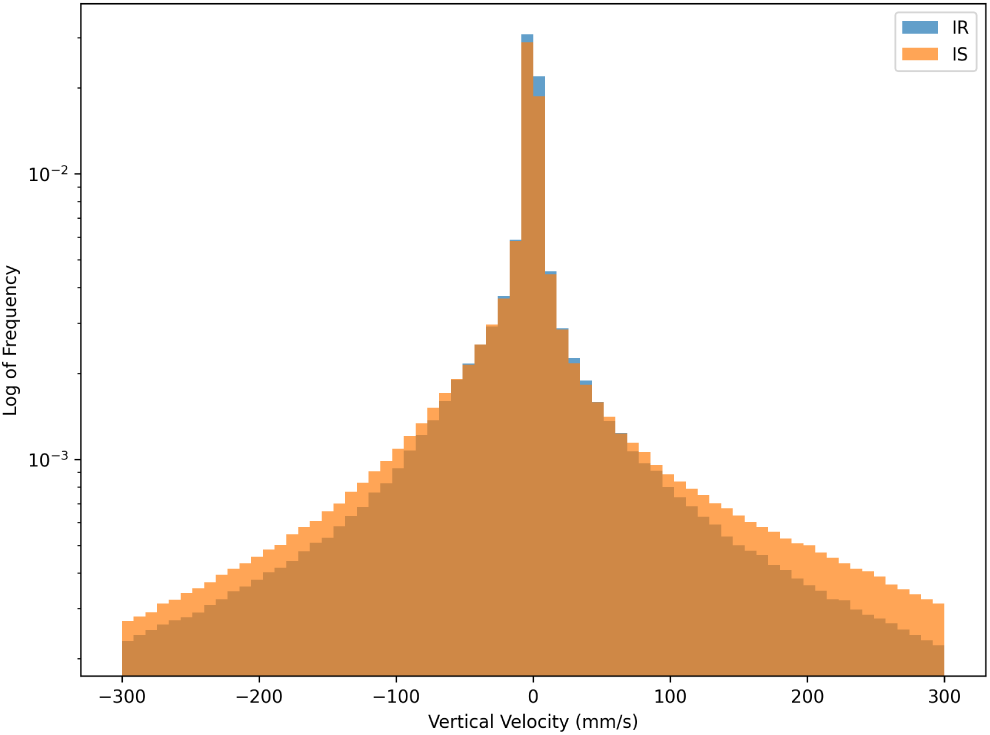
Histogram of vertical velocities for IR and IS mosquito trajectories. The velocity range is restricted within the range -300 mm/s to 300 mm/s to exclude outliers and extreme values. The plot uses logarithmic scaling for frequency.

Fractal dimension is another strong contributor in the separation between IR and IS, ranked 3rd in the SHAP bar plots. This feature is a measure of the linearity or complexity of the trajectory. Figure 7 shows the probability density histogram of trajectory segments for the XGBoost segment length (8 seconds) for each strain separately.

**Fig. 7:**
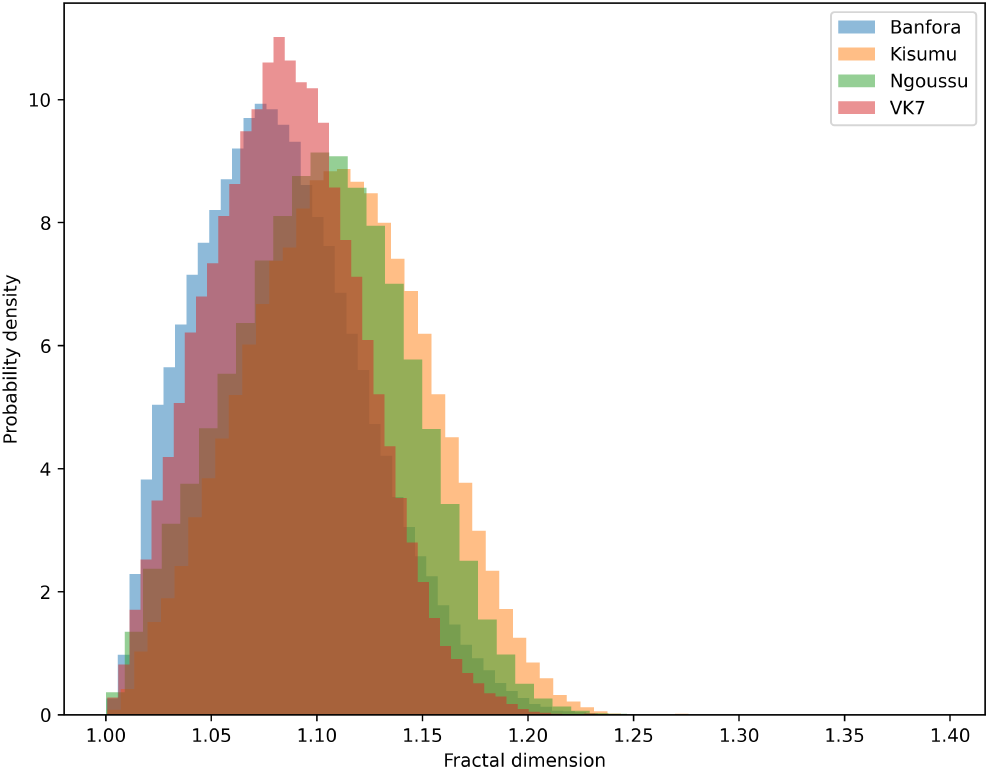
Probability density histogram of fractal dimension of trajectory segments, based on the XGBoost segment length (8 seconds), with each strain highlighted separately. The IR strains (Banfora in blue and VK7 in red) demonstrate lower fractal dimension values than the IS strains (Kisumu in orange and N’goussu in green). Interestingly, despite containing different strains, the inter-resistance classes show similarities in fractal dimension values. IR strains show more directed flight in comparison to IS.

### 2.3 Model Evaluation - Exploring Mosquito Strains

In addition to the analysis above, further target classifications were explored to assess the differences between the mosquito strains. Namely, whether in-class mosquito strains were separable and how different each strain was from one another.

#### 2.3.1 Classifying between Insecticide Susceptible (IS) strains

The differences within the insecticide susceptible class were explored, where a binary classification was attempted between the Kisumu and N’goussu strains using the XGBoost classifier. Table 4 displays the performance of this task. The confusion matrix is displayed in Figure 8 (A) which displays the percentage of the predictions over all folds. Additionally, the ROC curve is depicted in Figure 8 (B) featuring each folds’ curve.

**Fig. 8:**
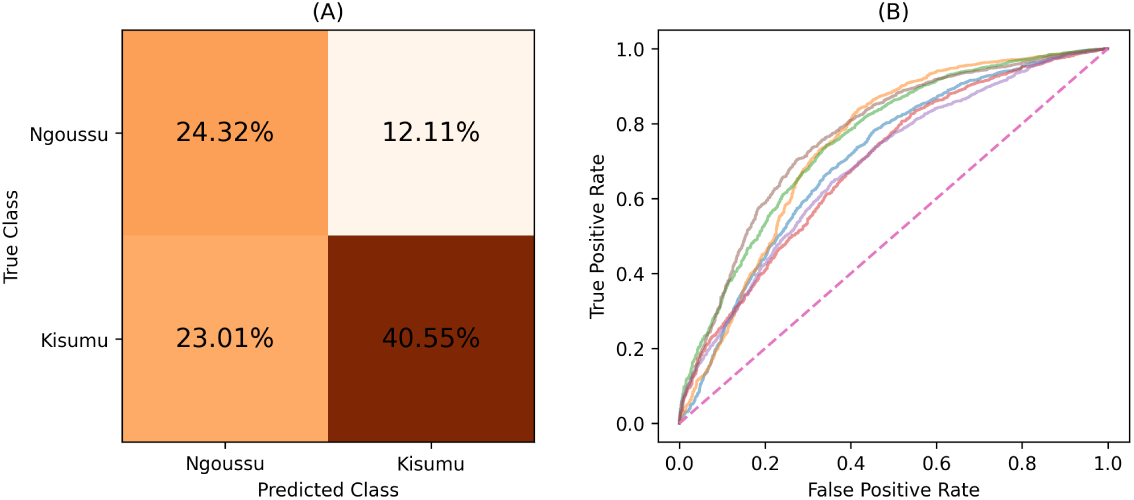
(A) Confusion matrix displaying the percentages of the predictions over all folds for the Kisumu vs N’goussu mosquito strains. (B) Figure displaying each folds’ ROC curve for the Kisumu vs N’goussu mosquito strains. Both graphs show results from independent test data.

**Table 4:** Performance of the XGBoost model applied to independent test data when classifying Kisumu and N’goussu mosquito strains.

| Model | Balanced Accuracy | ROC AUC score |
| --- | --- | --- |
| XGBoost | 0.655 (0.616 - 0.691) | 0.726 (0.688 - 0.768) |

#### 2.3.2 Classifying between Insecticide Resistant (IR) strains

Differences within the insecticide resistant class were also explored using binary classifiers, involving the Banfora and VK7 strains. Table 5 displays the performance of this task. The confusion matrix is displayed in Figure 9 (A) which shows the percentage of the predictions over all folds. Additionally, the ROC curve is in Figure 9 (B) featuring each folds’ curve.

**Fig. 9:**
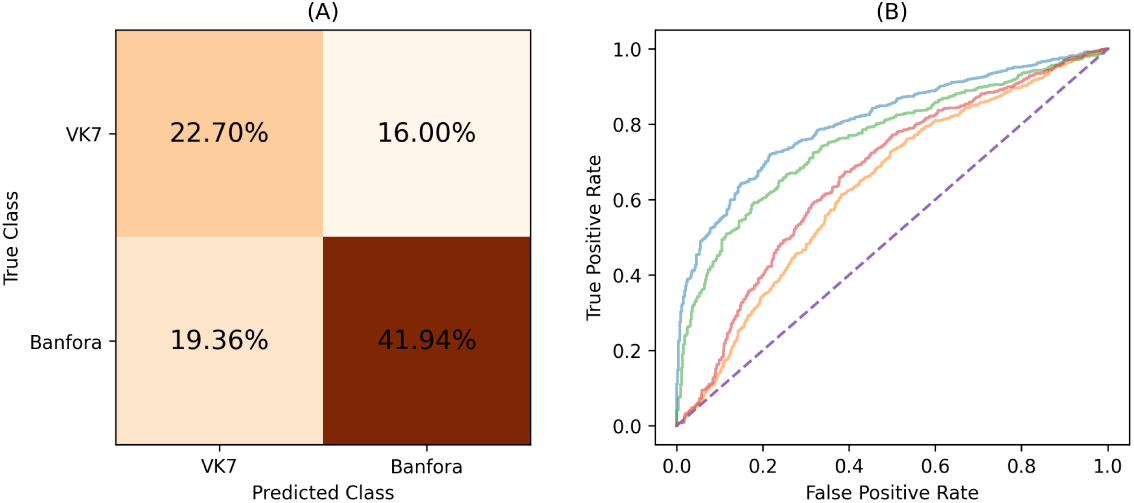
(A) Confusion matrix displaying the percentages of the predictions over all folds for the Banfora vs VK7 mosquito strains. (B) Figure displaying each folds’ ROC curve for the Banfora vs VK7 mosquito strains. Both graphs show results from independent test data.

**Table 5:** Performance of the XGBoost model applied to independent test data when classifying the Banfora and VK7 mosquito strains.

| Model | Balanced Accuracy | ROC AUC score |
| --- | --- | --- |
| XGBoost | 0.662 (0.606 - 0.725) | 0.717 (0.633 - 0.809) |

#### 2.3.3 Classifying between each strain

To assess the differences between each strain, we used a multiclass classification. The proposed pipeline remained mostly unchanged, however in this case the Mann Whitney U-test was performed pairwise for all strains and the multiclass XGBoost model was used. The results of the model are displayed in Table 6. The performance of each individual strain is also provided in Table 7, by calculating the model accuracy for each strain at each fold. The confusion matrix is displayed in Figure 10 which displays the sum of the predictions over all folds.

**Fig. 10:**
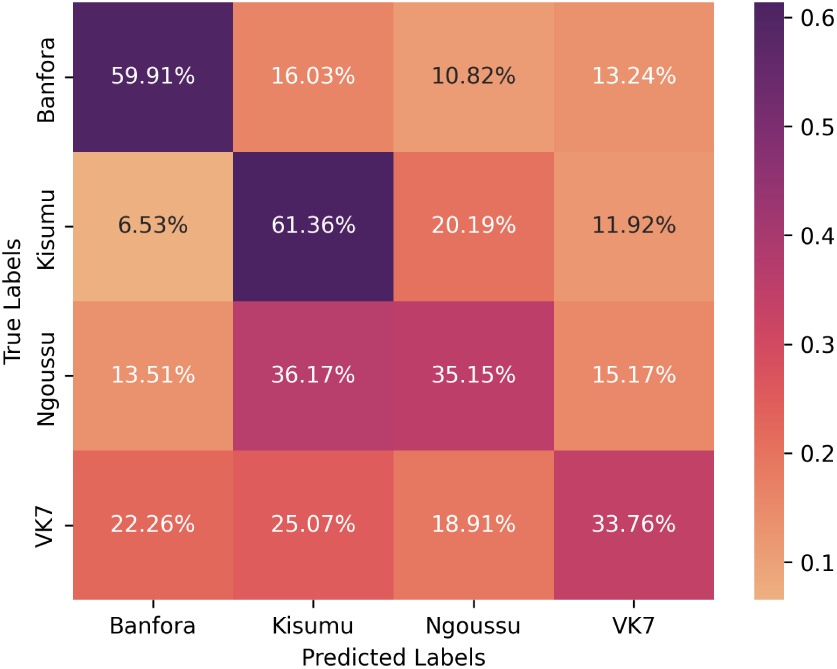
Confusion matrix displaying the performance of the multiclass classifier applied to independent test data. The colour and the value of each square represents the percentages of the predictions over all folds.

**Table 6:** Performance metrics for the multiclass classification model applied to independent test data with the average and the range (minimum and maximum) performance across all folds.

| Model | Balanced Accuracy (Training) | Balanced Accuracy (Testing) |
| --- | --- | --- |
| XGBoost | 0.928 (0.906 - 0.952) | 0.496 (0.454 - 0.553) |

**Table 7:** Multiclass model accuracy one-vs-all of each strain applied to independent test data with the average accuracy and the range provided in brackets.

| Mosquito Strain | Accuracy |
| --- | --- |
| Kisumu | 0.635 (0.515 - 0.845) |
| Banfora | 0.607 (0.422 - 0.813) |
| N'goussu | 0.368 (0.264 - 0.471) |
| VK7 | 0.354 (0.240 - 0.460) |

## 3 Discussion

ML models have been successful in classifying different mosquito strains based on flight trajectory features. The classification outcomes for the IR vs IS model with 2 mosquito strains in each class show the strongest performance using the non-linear XGBoost classifier with balanced accuracy and ROC AUC of 0.743 and 0.813 respectively. The model’s performance is consistent across all folds of the cross-validation process, see Figure 2. Uniquely, this was achieved without any data discriminating between resistant and susceptible mosquito responses to insecticide, but only on assumed differences between the two strains’ innate or inherent behaviours. Previously, responses of the same four IS and IR strains were investigated as they fed through ITNs and untreated (UT) net and the blood volume flow rate of the susceptible strain increased by 35% in presence of insecticide, but the resistant strain was already at the higher rate for both UT and ITNs (14). This supports the results presented here which demonstrate IR strains are behaving differently in the absence of an insecticide stimulus.

Whilst exploring the differences between IR and IS mosquito classes, the differences within the classes was examined. Within the susceptible class, the classifier identified notable differences between the Kisumu and N’goussu strains with an average balanced accuracy of 0.655 and a ROC AUC score of 0.726. Similarly, the classifier identified differences between the resistant strains, VK7 and Banfora, with an average balanced accuracy and ROC AUC score of 0.662 and 0.717, respectively. This illustrates that even though these strains are similar in their insecticide resistance, there are still identifiable differences between them. Yet, despite the differences within the classes, they still share common characteristics that enable the separation between IR and IS classes. A multiclass approach was also explored to identify distinctiveness of all four strains.

The performance of this approach was better than chance with an average balanced accuracy of 0.496. This may indicate that there are common behaviours across some or all mosquito strains. Nevertheless, an interesting feature of the multiclass results was the stronger separability of the Kisumu strain as well as the poor separability of VK7 (recall that features are evaluated over fixed duration segments and do not include activity levels). It is conjectured that the lower activity of VK7 may lead to its behaviour in short trajectory segments being encompassed by the other strains. In comparison, the stronger separability within the IR and IS classes may reflect genetic differences between the two IS strains and two IR strains. The susceptible strains are from different sides of the African continent (Kisumu originates from Kenya whereas N’gousso is from Cameroon) and were colonised more than 30 years apart. The two IR strains are both from southwest Burkina Faso, were both colonised in 2015 and both show similar phenotypic levels of pyrethroid resistance (10). However, the pyrethroid resistance in VK7 (2014) is largely conferred by target site resistance and elevated cytochrome P450 activity and whilst the Banfora strain has both of these resistance mechanisms there is also an indication that increased rates of respiration, and potentially changes in the microbiome also contribute to the resistance phenotype (15). The strongest performance is from the IR vs IS model utilising information from 2 genetically distinct IS and IR strains suggesting that there are common behavioural traits that evolve with IR strains that may associate with fitness benefits as described above.

The SHAPs analysis of the XGBoost IR vs IS model has identified the features which most strongly differentiate IR and IS behaviours. Interestingly, Figure 4 shows that there are non-negligible contributions for the vast majority of the features in the model – indicating that all features play a minor role in classification. IR behaviours are characterised by moving more slowly in the vertical direction than IS and hence expending less energy in flight. This also implies that on average an IR mosquito has lower momentum than IS and can change direction more quickly. For example, if the mosquito identifies blood meal cues such as CO2 or heat, IR behaviours indicate that it is flying more slowly and hence can adjust its trajectory more easily to follow those cues. The skewness of the distributions indicate that IR tend to fly down (negative velocities) more slowly than flying up in comparison to IS. In other words, IR fly more slowly when host seeking downwards, which may give more opportunity to detect and follow bloodmeal cues. Previous work identified that mosquitoes can detect a surface when some 30 to 45 mm away due to the induced velocity and pressure changes around the mosquito (16) observations that are consistent with experimental findings around a human baited bednet (12). This mechanism would lead to a reduction in velocity when flying downwards for all mosquitoes, however, the model here is comparing IR to IS strains. Hence, these results imply either an active response by the mosquito to the signals received when descending towards a surface or a more global behavioural shift or both. The skewed vertical velocity distribution also implies that IR fly upwards (positive velocity) more quickly; to escape predators. In making these interpretations it is important to note that in these experiments the mosquitoes have not blood fed – hence these are innate host seeking characteristics not influenced by taking on a blood meal and the considerable increase in mass that involves.

Horizontal-velocity features are largely similar to those for the vertical-velocity component in the standard deviation (ranked 9), 1st quartile (ranked 15) and 3rd quartile (ranked 26). IR behaviours have a narrower distribution of horizontal-velocities – implying that IR mosquitoes spend more time at slower speed than IS. The number of zero values and zero crossings (ranked 5) and number of local minima (ranked 8) introduce a further perspective. The number of zero crossings feature is more difficult to interpret as it includes the number of zero values of the horizontal-velocity within an 8 second segment. IR behaviours are associated through the model with low numbers of zero crossings or zero values, however, they have higher numbers of local minima than IS. This suggests that IR tracks are more consistent in direction compared to IS tracks. However, the higher number of local minima indicates a ‘jerky’ motion, characterised by frequent speeding up and slowing down. In the context of mosquito behaviour, a low amplitude ‘dither’ enables the mosquito to sample CO2 and thermal fields and maintain directed flight towards the highest concentration of these cues and hence a potential bloodmeal – see supplementary.

These differences in velocity features may be dependent on structure and performance of musculature, as well as a myriad of other parameters that may be optimal for these features in IS but not in IR. This could mean that there is a cost to being resistant, which leads to slower speeds and the inability to compete. There is evidence (17–19) which supports the notion that the mechanisms underpinning resistance may require additional energy, leading to a resource-based trade-off. This trade-off could explain the lower speeds and reduced variability in velocity features as observed in the SHAP analysis, Figure 5. These findings are further supported by raw trajectory data as shown in the histogram of positional velocity features, Figure 6. This lower variability in flight characteristics of IR strains could also be explained by the genetic bottleneck caused when selecting for resistance (e.g., fast blood-feeding). The resistant population may therefore be more genetically uniform showing less variation in features, e.g. vertical and horizontal velocity standard deviations, in comparison to IS mosquitoes. In the presence of ITNs, these characteristics become advantageous as they are selected for survival. However, in the absence of insecticide, this characteristic of IR could become a burden and hence would be lost.

Fractal dimension, ranked third in the model, measures the linearity or complexity of a trajectory. Values closer to 1 indicate more direct paths, while values approaching 2 signify complex paths with increased activity. SHAP analysis shows that IR has lower values than IS, approximately 1.04 for IR and 1.15 for IS. This trend aligns with many other features, consistently showing that IR tracks are more linear and direct compared to IS tracks. This potentially indicates that IR trajectories are more efficient or goal-oriented whilst IS tracks reflect more exploratory or less directed behaviour, see Figure 7, which illustrates that both IR strains, VK7 and Banfora, have distributions with similar lower values than the two IS strains, Kisumu and N’goussu.

Several features describe the shape of a trajectory. The centroid distance function (CDF) quantifies the deviations from the centroid, with a larger mean CDF indicating a wider or more variable track. SHAP analysis reveals that IR mosquitoes have a higher mean CDF (ranked 10), covering a larger area than IS, consistent with a larger convex hull (the 2D envelope of all positions in a segment) for IR. IR also has more local minima in CDF (ranked 22), suggesting less smooth and more oscillatory trajectories which may indicate bouncing behaviours. IS tends to stay closer to the centroid with fewer oscillations. The mean change in flight angle (ranked 13) is higher for IR, consistent with wider directional changes, while the 1st quartile (ranked 16) indicates a broader range for IR. IR has more zero-crossings in angle change (ranked 7), suggesting more consistent directional patterns supported by a higher number of local minima in CDF and higher tortuosity (ranked 37).

The analysis of angular velocity reveals that IR mosquitoes have a lower median angular velocity (ranked 23), suggesting smoother movements, but a higher mean angular velocity (ranked 25), indicating occasional sharp turns. This discrepancy suggests that while IR trajectories are generally smooth, they can still make sharp turns, likely due to their lower overall speed, which enhances manoeuvrability compared to IS. Curvature Scale Space (CSS) shows higher median CSS values for IS trajectories, implying sharper turns with smaller radii, consistent with IS exhibiting more complex, curved trajectories. In contrast, IR tracks are more direct with fewer pronounced turns, though occasional sharp turns may still occur.

In terms of accelerations, the vertical-axis component features generally support the discussion regarding preferential IR behaviours in vertical-velocity. The vertical-acceleration median (ranked 19) shows IR behaviours with an upwards bias whereas IS is zero to slightly downwards. Furthermore, the 3rd quartile of the vertical-acceleration distribution (ranked 21) is at higher positive values for IR than IS - supporting the ability of IR mosquitoes to generate stronger upward accelerations for escape. The significant horizontal-axis acceleration features are the 3rd quartile (ranked 20) which shows IS having larger values than IR and standard deviation (ranked 32) which shows IS having a broader distribution of values than IR. The horizontal-axis features are less clearly associated with fitness characteristics, and it is notable that these features are further down the ranking than for the vertical-axis and hence have lower influence on classification.

IR vs IS models were also determined using logistic regression and random forest classifiers, whilst the XGBoost classifier gave the best performance with average balanced accuracy 0.743 and ROC AUC 0.813. The worst performing classifier was logistic regression with an average balanced accuracy of 0.723 and a ROC AUC score of 0.788. This difference demonstrates that there are complex non-linear relationships across features when attempting to separate the classes. However, the performance of even the simpler linear models suggests that certain features exhibit linear separability. All models displayed stable results across folds with the range of performance metrics being consistently good (see Figure 2 (B) and (C) for the ROC curves which exhibit the expected slight variation between folds). This highlights that there are no abnormal trials in the dataset; the experiments were conducted in environmentally controlled laboratories minimising the variability of external effects. The small variation across folds also demonstrates the reliability and potentially high generalisation capability of this pipeline for classification tasks of trajectories.

There are some limitations to this study that should be addressed. One such limitation is that the behavioural differences identified may not be applicable to all IR and IS strains as only 4 mosquito strains were considered. Nevertheless, the classification task over all 4 strains revealed a large in-class diversity, and there were much stronger differences between IR and IS that were captured. Further analysis of the SHAP data could be conducted to examine consistency in trends across the folds. Future work should explore behaviours when exposed to a variety of ITN insecticides and more genetically distinct IR and IS mosquitoes. In summary, this paper demonstrates the ability of data driven machine learning classifiers to distinguish behaviours of pairs of IR and IS mosquito strains. SHAPs analysis identified that IR mosquitoes exhibit more directed flight together with a low amplitude ‘dither’ enabling the mosquito to sample the concentration of attractants and maintain flight towards the highest concentration of these cues and hence a potential bloodmeal. IR strains also fly slower on average vertically and horizontally – meaning they have less inertia and can change direction more easily in response to cues or threat.

## 4 Methods

Our approach for classifying the mosquito trajectories consists of several key steps. Mosquito trajectories are first split into segments using a sliding window method. These segments can then be used to generate features that describe the flight behaviour of the mosquitoes, such as the mean velocity, turning angle, and angular acceleration. These features are then provided to a machine learning model which is trained to classify segments as IS or IR. The classified segment predictions are combined using a voting method to form whole track predictions. The last stages include model evaluation and model interpretation using XAI techniques.

Throughout this work, a mosquito track was considered as a two-dimensional track or trajectory, *T*, that can be described as follows, where *x_i_* and *y_i_* correspond to the *i*-th position within a Cartesian coordinate system, and *t_i_*is time:

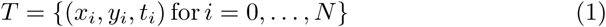

### 4.1 Dataset Description

The dataset used was generated within laboratory experiments at the Liverpool School of Tropical Medicine (LSTM), UK. In each experiment, three-to-five-day old unfed adult female mosquitoes from one of insecticide-susceptible (Kisumu and N’goussu) or insecticide-resistant (VK7 and Banfora) *An gambiae* strains were tracked around an untreated human-baited bednet for 2 hours (20). Further information about the strains used can be found in (21).

All experiments, also known as trials, were conducted between June 2019 and February 2020 using a custom built free-flight climate-controlled testing room (7 × 4.8 m in area, 2.5 m high). The experiments were performed in the afternoon to coincide with the ‘night’ phase of the mosquito’s circadian rhythm when they would be host-seeking in the wild. Mosquitoes were tracked using paired identical recording systems, where each recording system used one camera (12 MPixel Ximea CB120RG-CM with a 14 mm focal length lens), aligned with a single Fresnel lens (1400×1050 mm and 3 mm thick, 1.2 m focal length; NTKJ Co., Ltd, Japan) placed approximately 1210 mm away. The setup is telecentric and produces 2D data on mosquito flight. Further information on the experimental set up is outlined in (20) and the extraction of trajectories from video recordings is outlined in (22). Table 8 summarises the dataset used, where a strict limit on the track duration is set (above 1 second).

**Table 8:** Information on the trajectories of each mosquito strain.

| Strain | Insecticide-resistance Status | Number of Trials | Number of Tracks | Average track duration (s) | Average Velocity (mm/s) |
| --- | --- | --- | --- | --- | --- |
| Kisumu | Susceptible | 5 | 7631 | 22.39 (1.00-698.11) | 344.29 (5.20-2290.84) |
| N’goussu | Susceptible | 4 | 6125 | 21.40 (1.00-1010.03) | 453.78 (3.54-2602.18) |
| Banfora | Resistant | 4 | 7189 | 24.26 (1.00-475.42) | 389.57 (0.74-2478.15) |
| VK7 | Resistant | 4 | 5862 | 17.04 (1.00-457.77) | 429.54 (4.10-2102.89) |

The primary objective of this work is to explore differences between IR and IS mosquito flight behaviour, i.e. to consider Banfora and VK7 data together as IR and Kisumu and N’goussu as IS, as this will give more generic understanding and a larger dataset for the computational model. The outcomes of the IR vs IS model are analysed using the SHAPs XAI toolset (23) discussed in section 2.7. Further comparisons have been explored and ML model performances are reported: Kisumu vs N’goussu; Banfora vs VK7; and a multiclass model considering the 4 mosquito strains against each other.

### 4.2 Data Processing

The dataset used contains trajectories of varying durations, which can have an impact on the features generated and, in turn, affect the classification by the machine learning models. This variation in duration arises due to natural mosquito behaviour and the recording process. Small obstructions in the camera view, the mosquitoes sitting or walking on the bednet, as well as mosquitoes leaving the camera view, can break tracks into various lengths. Therefore, the trajectories in the dataset may only represent a portion of any mosquito’s flight path, and the reason there are different durations for each trajectory. To address this issue, trajectories were split into fixed duration segments using a moving window approach to unify track duration and eliminate duration bias for the machine learning model (24).

The window size and overlap between consecutive segments becomes a hyperparameter of the pipeline. Longer segments provide more information per segment but makes the model larger and increases the risk of overfitting. Larger overlap creates more segments per track improving per trajectory prediction but generates a lot of very similar segments again causing a risk of the model overfitting. To obtain a suitable window size and overlap, these parameters were optimised as described in section 2.8. The windowing approach ensures that each segment had the same duration, and thus the same amount of information, whilst excluding the length of the track as a feature (directly and indirectly) and removing the possibility for data leak.

As a result of the recording process, some positions within the mosquito tracks are missed. This can be due to mosquitoes flying in areas that were entirely obscured or encountering regions with poor contrast. To maintain the continuity of the tracks, linear interpolation was applied to fill in the gaps (25). However, in some instances, the tracks exhibited substantial gaps, introducing bias into the data. To address this issue and minimise its impact, we introduce a segment quality metric. It evaluates a track segment based on its information content, assigning higher scores to segments with more consecutively interpolated positions, as they contribute relatively little information. To establish a threshold for the metric, the mutual information of the tracks with the target variable (IR/IS status) at various thresholds of the segment quality metric was computed, where segments below this threshold were used, and the weighted average of the maximum thresholds that obtained the maximum mutual information for each trajectory feature was used as the final threshold. Track segments with segment quality scores larger than this threshold were removed. Various segment quality scores and thresholding techniques were assessed and are outlined in the supplementary information.

### 4.3 Feature Extraction

The extraction of meaningful features from the mosquito track segments is an essential step in the analysis of their flight patterns. The features can be broadly categorised into two types: shape descriptors and kinematic features. Shape descriptors capture the geometric characteristics of the tracks, such as the curvature of the path. On the other hand, kinematic features are based on time-dependent characteristics describing the movement dynamics, such as the speed, acceleration, and turning angle of the mosquito. After extraction of these characteristics, statistics (mean, standard deviation etc.) are computed for each feature over each segment and used as features for the model. As mentioned in the previous section, some of the positions within the tracks are interpolated so contain artificially generated positions. Feature statistics are calculated at positions containing real (not interpolated) data within a segment. The calculations for the features of flight are detailed in (24) for 3D trajectories and (26) for 2D trajectories and they are also provided in the supplementary.

### 4.4 Dataset Partitioning

The dataset was partitioned into two subsets: a tuning set and a modelling set. The tuning set was designated for feature selection and hyperparameters tuning. It comprises 2 trials for each strain, resulting in a total of 8 trials. Meanwhile, the modelling set, with a total of 9 trials, is dedicated to the actual machine learning model training and evaluation. Splitting the dataset by trials prevents data leakage and simulates a real-world scenario where an entire trial is tested. This division allows for robust model development and comprehensive evaluation of the machine learning models through cross-validation as well as elimination of data leaks.

### 4.5 Feature Selection

Having extracted meaningful features from trajectory segments, the Mann-Whitney U-test, a non-parametric statistical test, was employed to select relevant features using SciPy (27). To mitigate against group testing p-value inflation, a family-wise error rate (FWER) controlling procedure utilising Bonferroni correction was used. Features that demonstrated rejection of the null hypothesis at an FWER < 0.05 were selected. A Spearman correlation was calculated between all features. One feature from each pair of highly correlated features (ρ > 0.85) was removed from the set of selected features.

### 4.6 Classification Model

The objective of this study was to accurately differentiate between insecticide-susceptible and insecticide-resistant mosquito species using an ML classifier. To achieve this, a logistic regression, random forest, and XGBoost classification (28) models were evaluated. In order to generate whole track predictions, every track segment was classified independently and the mode of the segment binary predictions for each track was subsequently used as the prediction for the complete track. Prior to training the model, several processing steps were taken. Each feature was standardised through Z-score normalisation where the mean and standard deviation values were calculated from the training set. Then to balance the distribution of segments in each class, the training set was oversampled using SMOTE (Synthetic Minority Oversampling Technique) (29). These steps mitigate the models fitting to the imbalance of the dataset or to prioritise some features due to difference in the magnitude of their values.

### 4.7 Evaluation and Interpretation

The performance of the proposed framework was evaluated on the modelling dataset using a cross-validation approach. The dataset split in each iteration of this process is known as a fold, with each fold consisting of a different combination of two insecticide-resistant (IR - one from each IR strain) and two insecticide-susceptible (IS - one from each IS strain) trials in the training set, with the remainder of the trials used for model testing. Overall, there were 24 folds. When evaluating the performance of the training set, it should be noted that only those segments that were not generated via the oversampling technique (i.e., segments that constitute a genuine track) in the training set were used to derive the scores for the final track prediction.

Various performance metrics are calculated using Scikit-learn (28) including accuracy, balanced accuracy, ROC AUC (area under the receiver operator curve) score, PR AUC (area under the precision-recall curve) score, F1 score, precision score, recall score, Matthew Correlation Coefficient (MCC), Cohen kappa coefficient and log-loss score. The performance metrics are calculated on the whole track predictions where the arithmetic mean, and the minimum and maximum of performances across all folds is provided.

SHapley Additive ExPlanations (SHAP) (23) values are a powerful tool used to explain a model’s predictions. SHAP values represent the contribution of each feature to predictions made by a machine learning model. By calculating SHAP values for each feature, we can gain insights into which features have the strongest influence on the model’s output and how they interact with each other. In the context of studying insecticide resistance in mosquitoes, SHAP values were calculated for track segments and can help us understand the differences between IR and IS mosquito strains.

### 4.8 Hyperparameter Tuning

Each machine learning model has various parameters that require tuning, as well as the window size and overlap length of the windowing technique used to split tracks to segments. The parameters were tuned together in a cross-validated grid search approach attempting to maximise balanced accuracy. Tuning was conducted using two trials from each strain resulting in four IR trials and four IS trials. Each fold in the cross-validated grid search included each strain in the training set and test set. A full description of the parameter ranges and step sizes, as well as the full set of optimised hyperparameters identified for each model can be found in the supplementary.

