## Supplementary Information for "Discrimination of inherent characteristics of susceptible and resistant strains of *Anopheles gambiae* by explainable Artificial Intelligence Analysis of Flight Trajectories"

Y.M. Qureshi[1], V. Voloshin[1, 2], K. Gleave[3], H. Ranson[3], P.J. McCall[3],

J.A. Covington[1], C.E. Towers[1], D.P. Towers[1]

[1]School of Engineering, University of Warwick, Coventry, CV4 7AL, UK

[2]School of Biological and Behavioural Sciences, Queen Mary University of London, E1 4NS, UK

[3]Vector Biology Department, Liverpool School of Tropical Medicine, Pembroke Place, Liverpool, L3 5QA, UK

Corresponding Author:

 (YMQ)

**Feature Calculations**

1. Velocity, acceleration and Jerk

These are calculated using second-order central finite difference methods. The signed values are used – where in the case of velocity, a positive velocity indicates motion from head-to-toe of the human bait within the experiment. The axial values are also utilised.

2. Angle of Flight

The angle of flight, $\alpha_i$, can be calculated using the dot product of two vectors. It describes the relative change in direction between positions.

$$v_i = \begin{bmatrix} x_i - x_{i-1} \\ y_i - y_{i-1} \end{bmatrix}$$

$$v_{i+1} = \begin{bmatrix} x_{i+1} - x_i \\ y_{i+1} - y_i \end{bmatrix}$$

$$\alpha_i = \arccos\left(\frac{v_i \cdot v_{i+1}}{\sqrt{(v_i \cdot v_i)(v_{i+1} \cdot v_{i+1})}}\right)$$

3. Angular Velocity and Angular Acceleration

The angular velocity and angular acceleration are both calculated using second-order finite difference methods. The arctangent function is used to calculate the angle between positions, then the change in angle between consecutive positions is computed. Angular velocity is calculated by dividing this change in angle by the corresponding time difference.

4. Orthogonal Components of Velocity

This is a group of features that describe the tendency of a movement to head in a given direction. The persistence velocity describes the tendency to head tangential to the trajectory while the turning velocity describes the tendency to head normal to the trajectory. It involves converting from the Cartesian to the circular coordinate system:

$$\rho_i = \sqrt{(x_{i+1} - x_i)^2 + (y_{i+1} - y_i)^2}$$

$$\theta_i = \arctan\left(\frac{y_{i+1} - y_i}{x_{i+1} - x_i}\right)$$

The change in $\theta$ and instantaneous velocity is calculated:

$$\Theta_i = |\theta_{i+1} - \theta_i|$$

$$v_i = \frac{\rho_i}{t_{i+1} - t_i}$$

Then converting back to the Cartesian coordinate system:

$$P_i = v_i \cos \Theta$$

$$T_i = v_i \sin \Theta$$

Where $P_i$ and $T_i$ are the persistence and turning velocities, respectively.

5. Tortuosity

This is the ratio of the actual distance travelled and the shortest distance between the start and end positions.

$$S = \frac{\sum_{i=0}^{N} \sqrt{(x_{i+1} - x_i)^2 + (y_{i+1} - y_i)^2}}{\sqrt{(x_N - x_0)^2 + (y_N - y_0)^2}}$$

6. Convex Hull

The convex hull is the set of points that forms the smallest possible convex polygon that bounds all the points of a track. From the polygon that is formed, the area and perimeter is extracted and used as features.

7. Centroid Distance Function

This calculated the distances of each position in a track to the centre point of the track.

$$x_c = \frac{1}{N} \sum_{i=0}^{N} x_i$$

$$y_c = \frac{1}{N} \sum_{i=0}^{N} y_i$$

$$C_i = \sqrt{(x_i - x_c)^2 + (y_i - y_c)^2}$$

8. Curvature

Curvature is a measure that defines the deviation of a trajectory from a straight line. It is calculated as:

$$k_i = \frac{\dot{x}_i \ddot{y}_i - \dot{y}_i \ddot{x}_i}{\left(\dot{x}_i^{\,2} + \dot{y}_i^{\,2}\right)^{\frac{3}{2}}}$$

9. Curvature Scale Space

This measures the inflection points in a trajectory at difference scales and is robust to affine transformations. To compute the curvature at varying levels, the trajectory is convolved with a 1D gaussian kernel of width $\sigma$. The value of $\sigma$ is varies, and the zero-crossings locations in the curvature are recorded. The locations of zero-crossings at different $\sigma$ levels can be plotted as a CSS image. A 1D signal is generated by taking column maximums from this CSS image.

10. Fractal Dimension

Fractal dimension is a measure of the complexity of a trajectory. Values around 1 describe a linear path with a shape similar to a straight line, and values around 2 indicate convoluted movement with a shape similar to a plane. To compute fractal dimension, $d$, the following equations are used:

$$d = \frac{\log n}{\log \frac{1}{S}}$$

Where $n$ is the number of minature pieces in the path, $S$ is the scaling factor, and $d$ is the fractal dimension. For a trajectory, the values of $n$ and $S$ are defined as follows:

$$n = \sum_{i=0}^{N} \sqrt{(x_{i+1} - x_i)^2 + (y_{i+1} - y_i)^2}$$

$$S = \frac{1}{\sqrt{(\max x - \min x)^2 + (\max y - \min y)^2}}$$

**Penalty functions and thresholds assessed.**

To remove tracks with substantial gaps (many missing positions), various methods were assessed. These include:

1. Removing tracks with percentage of interpolated positions

If the track contains greater than or equal to n% of interpolated positions, it is removed. A few values were tested including 25%, 50%, 75%.

2. Removing tracks with large number of consecutive positions

If the tracks contains greater than or equal to a number of interpolated positions, it is removed. The threshold for this was assessed by the percentage of the whole track, where 25%, 50% and 75% were assessed.

3. Removing tracks with penalty score larger than a threshold

This is where we developed a penalty function to score a track on its information content. Let $segment$ be a list of length $N$ where each element in the list is either 1 or 0, indicating a real position (1) or an interpolated position (0) for a segment. The penalty function $P(segment, n, m)$ is defined as:

$$P(segment, n, m) = \frac{1}{N}\sum_{i=0}^{N}\begin{cases} n \cdot m^{c_i}, & \text{if } x_i = 0 \\ 0, & \text{if } x_i = 1 \end{cases}$$

Where $c_i$ is the length of the consecutive artificial positions starting from position $i$ until a real position is encountered. If $x_i = 1$, $c_i$ is reset to 0. Various values of $n$ and $m$ were assessed with $n = 1$ and $m = 1.05$ being used for the final model.

After each track is provided with a score, we can identify a threshold. A few different methods were tested. These include:

a. Mutual Information-based Threshold

To obtain a threshold for the penalty score, a threshold was determined by computing the mutual information at various score thresholds. Then the weighted average threshold is obtained based on the maximum mutual information and its corresponding threshold for each feature. This weighted average is the final score threshold.

b. Mutual information and Gradient-based Threshold

To obtain a threshold for the penalty score, the threshold is identified by computing the mutual information at various score thresholds, but then using the gradient to obtain a threshold for each feature. The gradient is the rate of change in mutual information, where the threshold is identified where the rate of change drops below a predefined value. Finally, the weighted average of these identified thresholds based on their associated mutual information values is computed to return the final threshold.

**Comparison to Dither**

The SHAP plots indicate that higher number of local minima in horizontal-velocity features tend towards IR. This suggests that IR tracks have jerky motion with frequent speeding up and slowing down. This behaviour is analogous to the deliberate use of 'dither' in engineering control systems where it has been used successfully to minimise vibration noise (Kropp et al "The application of dither to mitigate curve squeal" (2021) Journal of Sound and Vibration, 514, art. no. 116433) and produce low vibration control of magnetic levitation vehicles (Tomono et al "Controllability improvement of zero-power magnetically levitated system by dither", (1998) IEEE Annual Power Electronics Specialists Conference, 1, art. no. 701958, pp. 588 – 593). This low amplitude 'dither' in flight may allow the mosquito to be aware of its surroundings by sampling $CO_2$ and thermal fields. This could then lead to directed flight towards these cues which indicates a potential bloodmeal.

Table 1. Hypertuning parameter ranges for XGBoost

| Parameter | Parameter values |
|---|---|
| Window size | 0.5, 1, 1.5, 2, 2.5, 3, 3.5, 4, 4.5, 5, 5.5, 6, 6.5, 7, 7.5, 8, 8.5, 9, 9.5 |
| Window overlap | 0.5, 1, 1.5, 2, 2.5, 3, 3.5, 4, 4.5, 5, 5.5, 6, 6.5, 7, 7.5, 8, 8.5, 9 |
| Learning rate | 0.01, 0.1, 0.3 |
| N estimators | 50, 100, 150, 200, 250 |
| Max depth | 3, 5, 7 |
| Subsample | 0.5, 0.7, 0.9 |
| Colsample bytree | 0.5, 0.7, 0.9 |
| Reg alpha | 0, 0.01, 0.1 |
| Reg lambda | 0, 0.01, 0.1 |
| Min child weight | 1, 5, 15 |

Table 2. Hypertuning parameter ranges for random forests

| Parameter | Parameter values |
|---|---|
| Window size | 0.5, 1, 1.5, 2, 2.5, 3, 3.5, 4, 4.5, 5, 5.5, 6, 6.5, 7, 7.5, 8, 8.5, 9, 9.5 |
| Window overlap | 0.5, 1, 1.5, 2, 2.5, 3, 3.5, 4, 4.5, 5, 5.5, 6, 6.5, 7, 7.5, 8, 8.5, 9 |
| N estimators | 100, 200, 300, 400 |
| Criterion | Entropy, gini |
| Max depth | 5, 10, 15 |
| Min samples split | 2, 3, 4 |
| Min samples leaf | 1, 2, 3 |
| Max depth | Log2, sqrt |

| | |
|---|---|
| Bootstrap | True, False |

Table 3. Hypertuning parameter ranges for logistic regression

| Parameter | Parameter values |
|---|---|
| Window size | 0.5, 1, 1.5, 2, 2.5, 3, 3.5, 4, 4.5, 5, 5.5, 6, 6.5, 7, 7.5, 8, 8.5, 9, 9.5 |
| Window overlap | 0.5, 1, 1.5, 2, 2.5, 3, 3.5, 4, 4.5, 5, 5.5, 6, 6.5, 7, 7.5, 8, 8.5, 9 |
| Penalty | L1, L2, None |
| c | 0.01, 0.1, 1, 10, 100 |
| Solver | lbfgs, liblinear, newton-cg, sag, saga |
| Max iteration | 50, 100, 150 |

Table 4. Hypertuning parameter ranges for Kisumu vs Ngoussu – XGBoost

| Parameter | Parameter values |
|---|---|
| Window size | 0.5, 1, 1.5, 2, 2.5, 3, 3.5, 4, 4.5, 5, 5.5, 6, 6.5, 7, 7.5, 8, 8.5, 9, 9.5 |
| Window overlap | 0.5, 1, 1.5, 2, 2.5, 3, 3.5, 4, 4.5, 5, 5.5, 6, 6.5, 7, 7.5, 8, 8.5, 9 |
| Learning rate | 0.01, 0.1, 0.3 |
| N estimators | 100, 150, 200 |
| Max depth | 3, 5, 7 |
| Subsample | 0.5, 0.7, 0.9 |
| Colsample bytree | 0.5, 0.7, 0.9 |
| Reg alpha | 0, 0.01, 0.1 |
| Reg lambda | 0, 0.01, 0.1 |

| | |
|---|---|
| Min child weight | 1, 5, 15 |

Table 5. Hypertuning parameter ranges for Banfora vs VK7 – XGBoost

| Parameter | Parameter values |
|---|---|
| Window size | 0.5, 1, 1.5, 2, 2.5, 3, 3.5, 4, 4.5, 5, 5.5, 6, 6.5, 7, 7.5, 8, 8.5, 9, 9.5 |
| Window overlap | 0.5, 1, 1.5, 2, 2.5, 3, 3.5, 4, 4.5, 5, 5.5, 6, 6.5, 7, 7.5, 8, 8.5, 9 |
| Learning rate | 0.01, 0.1, 0.3 |
| N estimators | 100, 150, 200 |
| Max depth | 3, 5, 7 |
| Subsample | 0.5, 0.7, 0.9 |
| Colsample bytree | 0.5, 0.7, 0.9 |
| Reg alpha | 0, 0.01, 0.1 |
| Reg lambda | 0, 0.01, 0.1 |
| Min child weight | 1, 5, 15 |

Table 6. Hypertuning parameter ranges for Multiclass XGBoost

| Parameter | Parameter values |
|---|---|
| Window size | 0.5, 1, 1.5, 2, 2.5, 3, 3.5, 4, 4.5, 5, 5.5, 6, 6.5, 7, 7.5, 8, |
| Window overlap | 0.5, 1, 1.5, 2, 2.5, 3, 3.5, 4, 4.5, 5, 5.5, 6, 6.5, 7, 7.5 |
| Learning rate | 0.01, 0.1, 0.3 |
| N estimators | 100, 150, 200 |
| Max depth | 3, 5, 7 |
| Subsample | 0.5, 0.7, 0.9 |

| | |
|---|---|
| Colsample bytree | 0.5, 0.7, 0.9 |
| Reg alpha | 0, 0.01, 0.1 |
| Reg lambda | 0, 0.01, 0.1 |
| Min child weight | 1, 5, 15 |

Table 7. Final hyperparameters for XGBoost model

| Parameter | Parameter values |
|---|---|
| Window size | 8 |
| Window overlap | 7.5 |
| Learning rate | 0.3 |
| N estimators | 250 |
| Max depth | 5 |
| Subsample | 0.5 |
| Colsample bytree | 0.9 |
| Reg alpha | 0 |
| Reg lambda | 0 |
| Min child weight | 15 |

Table 2. Final hyperparameters for random forests

| Parameter | Parameter values |
|---|---|
| Window size | 7.5 |
| Window overlap | 6.5 |
| N estimators | 300 |
| Criterion | Entropy |
| Max depth | 15 |
| Min samples split | 3 |
| Min samples leaf | 1 |
| Max depth | sqrt |
| Bootstrap | False |

Table 3. Final hyperparameters for logistic regression

| Parameter | Parameter values |
|---|---|
| Window size | 9.5 |
| Window overlap | 9 |
| Penalty | L2 |
| c | 0.01 |
| Solver | Saga |
| Max iteration | 50 |

Table 4. Final hyperparameters for Kisumu vs Ngoussu – XGBoost

| Parameter | Parameter values |
|---|---|
| Window size | 1.5 |
| Window overlap | 0.5 |
| Learning rate | 0.1 |
| N estimators | 200 |
| Max depth | 5 |
| Subsample | 0.5 |
| Colsample bytree | 0.7 |
| Reg alpha | 0.01 |
| Reg lambda | 0.1 |
| Min child weight | 15 |

Table 5. Final hyperparameters for Banfora vs VK7 – XGBoost

| Parameter | Parameter values |
|---|---|
| Window size | 6 |
| Window overlap | 5.5 |

| | |
|---|---|
| Learning rate | 0.1 |
| N estimators | 200 |
| Max depth | 3 |
| Subsample | 0.5 |
| Colsample bytree | 0.9 |
| Reg alpha | 0.01 |
| Reg lambda | 0.1 |
| Min child weight | 15 |

Table 6. Final hyperparameters for Multiclass XGBoost

| Parameter | Parameter values |
|---|---|
| Window size | 6 |
| Window overlap | 5.5 |
| Learning rate | 0.3 |
| N estimators | 200 |
| Max depth | 5 |
| Subsample | 0.5 |
| Colsample bytree | 0.5 |
| Reg alpha | 0.1 |
| Reg lambda | 0.1 |
| Min child weight | 5 |

Figure 1. Confusion matrix for insecticide resistant vs insecticide susceptible task with sum of tracks across all folds, where (A) is the logistic regression model, (B) is the random forest model and (C) is the XGBoost model.

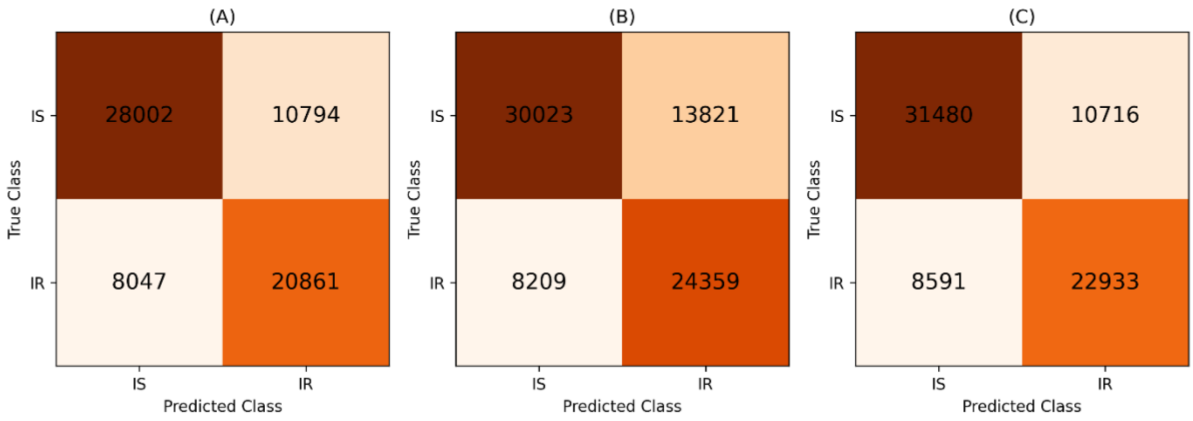


Figure 2. Confusion matrix Kisumu vs N'goussu task with sum of tracks across all folds.

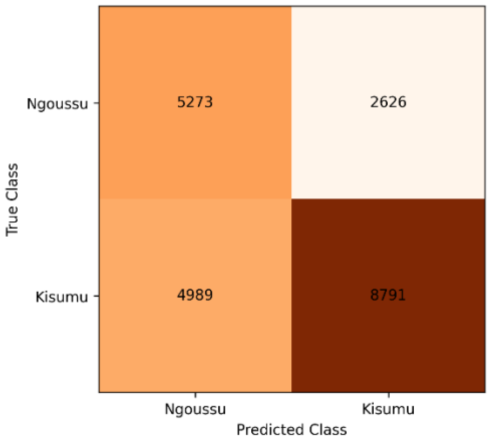


Figure 3. Confusion matrix Banfora vs VK7 task with sum of tracks across all folds.

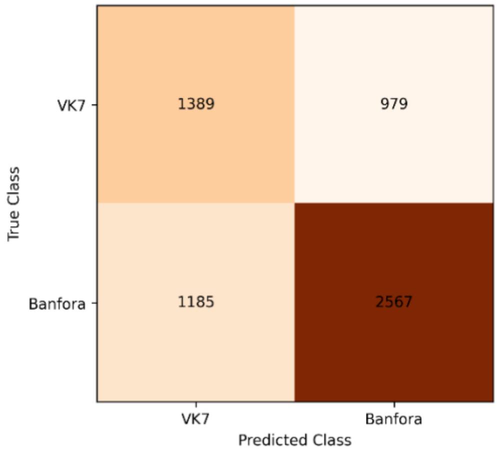
